# Chronic Stress Alters Astrocyte Morphology in Mouse Prefrontal Cortex

**DOI:** 10.1101/2021.02.23.432559

**Authors:** Sierra A. Codeluppi, Dipashree Chatterjee, Thomas D. Prevot, Yashika Bansal, Keith A. Misquitta, Etienne Sibille, Mounira Banasr

## Abstract

**Background:** Neuromorphological changes are consistently reported in the prefrontal cortex (PFC) of patients with stress-related disorders and in rodent stress models, but the effects of stress on astrocyte morphology and potential link to behavioral deficits are relatively unknown.

**Methods:** To answer these questions, transgenic mice expressing green fluorescent protein (GFP) under the glial fibrillary acid protein (GFAP) promotor were subjected to 7, 21 or 35 days of chronic restraint stress (CRS). CRS induced behavioral effects on anhedonia- and anxiety-like behaviors were measured using the sucrose intake and the PhenoTyper tests, respectively. PFC GFP+ or GFAP+ cells morphology was assessed using Sholl analysis and associations with behavior were determined using correlation analysis.

**Results:** CRS-exposed male and female mice displayed anxiety-like behavior at 7, 21 and 35 days and anhedonia-like behavior at 35 days. Analysis of GFAP+ cell morphology revealed significant atrophy of distal processes following 21 and 35 days of CRS. CRS induced similar decreases in intersections at distal radii for GFP+ cells, accompanied by increased proximal processes. In males, the number of intersections at the most distal radius step significantly correlated with anhedonia-like behavior (r=0.622, p<0.05) for GFP+ cells and with behavioral emotionality calculated by z-scoring all behavioral measured deficits (r=-0.667, p<0.05). Similar but not significant correlations were observed in females. No correlation between GFP+ cell atrophy with anxiety-like behavior was found.

**Conclusion:** Chronic stress exposure induces a progressive atrophy of cortical astroglial cells, potentially contributing to maladaptive neuroplastic and behavioral changes associated with stress-related disorders.

## Introduction

Allostatic response plays a significant role in an organism ability to cope with stressful life events by meeting the environmental challenge demands and return to a homeostatic state [1, 2]. However, following repeated stress exposure the increase in allostatic load and the homeostatic state can no longer be maintained. This is associated with maladaptive responses within limbic brain regions, linked with several CNS stress-related disorders including major depressive disorder (MDD) [1-4]. One particular brain region is the prefrontal cortex (PFC), which is important in the top-down control of stress response and has been implicated in cognition and emotion regulation [2]. The PFC undergoes various neuroplastic changes in response to chronic stress exposure that include morphological reorganization such as altered dendrite arborization, dendrites retraction and synaptic loss [1, 5]. These effects are reversible when stress exposure is interrupted [6], but if chronic stress persists, it can lead to morphological changes similar to those observed in MDD. These changes include PFC volume reductions [7-9] and reduced size of PFC neurons and glial cell numbers [10, 11]. Several brain imaging studies have reported that patients with MDD showed volumetric changes in the PFC [12-14]. Although neuronal loss with MDD is controversial [15], studies have consistently reported astroglial loss [16, 17] and reduced synapse number and synaptic markers [18-20].

Astroglia cells make up ∼40% of the cells in the human brain and are numerous in the PFC [21]. Astrocyte morphology is classically described as a star-like cell structure with the center being the soma and distinctive long processes that protrude outward. Each of these processes display smaller branching processes of varying lengths and sizes [22]. Astrocytes are crucial for proper function and CNS plasticity since they are important regulators of synaptic connectivity, synaptogenesis, and maintenance of synaptic function. In addition to providing energy, nutrients and trophic factors to neurons as well as contributing to recycling neurotransmitters, astrocytes contribute to the fabrication and maintenance of the extracellular matrix responsible for synapse morphology [23-27].

Astroglia are likely key mediators in the synaptic dysfunction implicated in multiple pathologies of mood disorders [28-34]. Post-mortem and preclinical studies consistently report reduced density and number of GFAP+ astroglial cells [16, 35]. Studies also reported decreased GFAP protein and mRNA levels in key brain regions of patients with mood disorders [35, 36], reductions of GFAP+ cells in the PFC of MDD patients [11], and reductions of various specific markers of astroglial cell function [36]. In rodent models of chronic stress, fewer GFAP+ cells are reported in the hippocampus [37], amygdala and subregions of the PFC [16]. Previous work from our group demonstrated that animals subjected to chronic stress show reduced levels of GFAP mRNA expression, fewer GFAP+ cells in the prelimbic cortex and reduced glial metabolism in the PFC [38, 39]. Interestingly, these changes appear selective to GFAP+ astroglia since cortical S100β+ cell density remains unaffected [40], suggesting that GFAP+ astroglia population is especially susceptible to chronic stress. Cortical GFAP+ astroglia reduction was directly involved in the expression of depressive-like behaviors. Indeed, we previously demonstrated that local infusion of a gliotoxin producing ablation of GFAP+ cells in the PFC induced anxiety- and anhedonia-like deficits similar to those seen following chronic stress [38]. Similar PFC astroglia ablation impairs cognitive flexibility [41], promotes ethanol intake [42] and prevents the antidepressant action of deep brain stimulation [43].

Despite numerous studies identifying reduced astroglial density, little is known on the effects of chronic stress on astroglial morphology in the PFC. In this study, we aimed to conduct comprehensive analyses of chronic stress effects on PFC GFAP+ astroglial morphology using the chronic restraint stress (CRS) model and Sholl analysis [6, 44-48]. We first assessed the behavioral consequences of 7, 21, or 35 days of CRS in transgenic mice expressing green fluorescent protein (GFP) under the GFAP promotor. We then conducted a Sholl analysis on GFAP astroglia immunolabelled with either GFAP (primarily staining major processes) or GFP (allowing staining of additional small processes). In the investigation of neuronal cell morphology, Sholl analysis and the quantification of the number of intersections is the classical method used to assess branching complexity [49]. Since neuronal morphology is relatively homogeneous within a brain region, this method is well suited for studying morphological complexity of neurons. Several studies have attempted to apply the same principles to astrocytes [50, 51] but were unable to detect chronic stress effects [52], most likely because astrocytes are highly heterogeneous in size and wildly different in morphology [53, 54]. In this study, our goal was to adapt the Sholl analysis method to astrocytes and assess astroglial morphology after stress and potential correlation with behavioral deficits.

## Methods

### Animals

Animals were housed under normal conditions with an artificial 12hr light/dark cycle and *ad libitum* access to food and water (except when deprived for testing). All animals were single housed starting the week before the beginning of the stress experiments, for individual monitoring of behavioral assessments. All procedures and experiments followed guidelines of the Canadian Council on Animal Care (CCAC) and were approved by Centre for Addiction and Mental Health (CAMH) animal care committee (ACC). We used FVB/N-Tg (GFAPGFP)14Mes/J transgenic mice (Jackson Laboratories, stock #003257) backcrossed with wild type (WT) C57BL/6 mice, for at least seven generations. Heterozygous GFAP-GFP mice (n=32) were 8-16 weeks of age at the beginning of the experiment.

### Chronic Restraint Stress (CRS)

Four groups (n=8/group, 50% females) were used in this study i.e n=4/sex/group (**Figure 1a**): control group (CRS 0) and animals subjected to 7 (CRS 7), 21 (CRS 21), or 35 days of CRS (CRS 35). During CRS, animals were placed in Falcon® Tubes with holes at the bottom and the cap to allow for air flow, 1 hour, twice a day (2 hours apart) between 11:00 am to 4:00 pm. Control animals were handled daily. Similar CRS procedures were previously used to assess effects of chronic stress on neuronal morphology [55-57].

**Figure 1:**
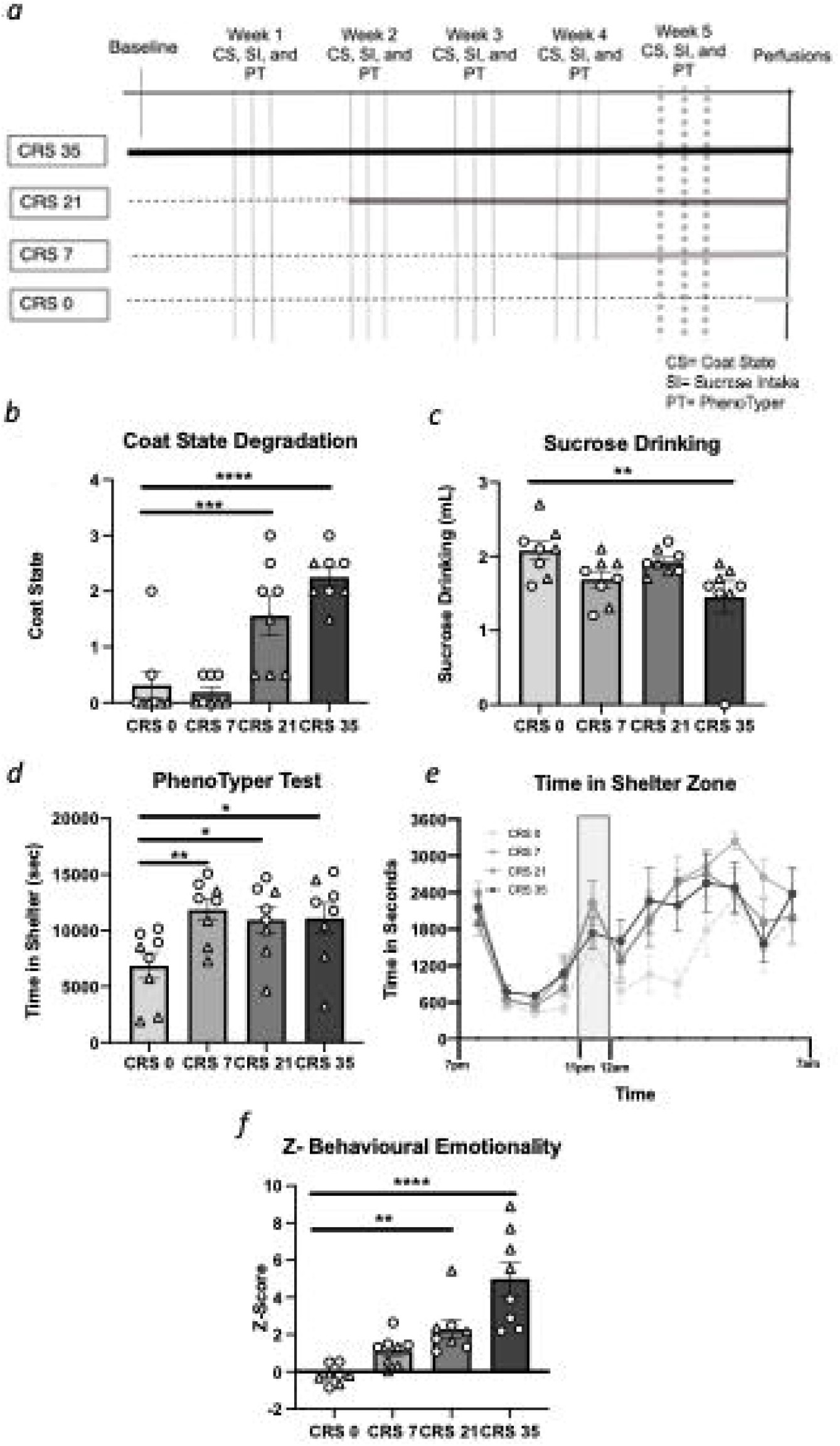
Chronic restraint stress (CRS) effects on behavioral emotionality at experimental week *5*. **(a)** Timeline of CRS and behavioral testing. Behavioral performances measured throughout the chronic stress exposure can be found in **Supplementary Materials**, this figure only illustrates the results obtained on week 5. CRS increases coat state degradation **(b)** and reduces sucrose intake **(c)**. In the PhenoTyper test, CRS increases time in the shelter measured either for 5 hours after the light challenge **(d)** or hourly **(e)**. CRS increases the overall z-score of behavioral emotionality **(f)**. All data is presented as mean ± s.e.m.*\* p<0*.*05*, ** *p<0*.*01, *** p<0*.*001* and *\*\*\*\* p<0*.*0001* compared to control group. Females are denoted with a triangle and males with a circle.

### Longitudinal behavioral Assessments

To assess the progression of chronic stress, animals were tested at the behavioral level weekly; including assessment of coat state, sucrose intake, and time spent in the shelter during the PhenoTyper test (**Figure 1a**). The experimenter was blinded to the experimental group assignments.

### Coat State Assessment

Coat state assessment was performed as described by Yalcin et al. [58]. Each week 18 hours after the restraint stress session, seven body parts were assessed for coat quality (i.e., the head, neck, dorsal coat, ventral coat, tail, forepaws, and hind paws). Each body part was given a score of 0, 0.5, or 1 from well-kept to unkempt. Sum of all scores determined the deterioration score of the animal’s coat state.

### Sucrose consumption test

During the first week animals were habituated for 48 hours to 1 % sucrose solution. After habituation, mice were fluid deprived for 16 hours overnight which was followed by 1 hour of sucrose consumption test. After the sucrose consumption test, animals were returned back on tap water for 24 hours. After 24 hours, animals were deprived of water for 16 hours and then, water intake was measured for 1 hour. Every following week, the animals are only re-habituated to sucrose for 24h before the test. Sucrose and water consumption tests were performed each week 18 hours after the restraint stress session.

### PhenoTyper Test

The PhenoTyper test is a consistent and reliable test that measure conflict anxiety-like behavior in mice [59-61]. The test uses the PhenoTyper® (Noldus, Leesburg, VA, USA) apparatus, a 30 cm^2^ home cage-like arena with a designated shelter zone and food zone. Each week, animals are placed in the PhenoTyper (minimum of 3 hours after the restraint session and recorded during their dark cycle (19:00 to 7:00). A spotlight placed above the food zone is programmed to turn on at 23:00 for 1 hour. The light challenge causes mice to hide in the shelter zone; behavior exacerbated by chronic stress [61]. The continued avoidance of the food zone in favor to shelter is calculated for each animal as the time spent in the shelter zone for the five hours after the light challenge (from 12 am to 5am).

### Z-Emotionality Score

We used an integrative z-scoring method to assess the consistency of behavior across tests and compiling behavioral readouts into one score of behavioral emotionality for each animal. The z-behavioral emotionality score is calculated as a function of the standard deviation and average of the control group for each sex and as an average of performances of all behavioral tests [62].

### Mouse Euthanasia and Perfusion

Animals were anesthetized 20 hours after the last stressor with avertin (125mg/kg, i.p.) and perfused intracardially with 4% paraformaldehyde (PFA) using a Pharmacia pump (Cole-Parmer, Montreal, Canada). Brains were post-fixed in 4% PFA overnight at 4°C, cryoprotected in a 30% sucrose solution (48h at 4°C) and frozen on dry ice before being stored at -80°C. Using a cryostat (Leica CM1950, Illinois, USA), 40μm-thick sections containing the PFC region (anterioposteriority 2.3-1.4mm from Bregma) were cut and placed in cryoprotectant (30% sucrose, 1% PVP-40, 30% ethylene glycol) for storage at -20°C.

### Immunohistochemistry

Free-floating sections were washed in 0.3% PBS (phosphate buffer saline)-TritonX (PBS-T), then incubated with 3% H_2_O_2_ in PBS for 20 mins at room temperature. Sections were then incubated for 1h in 10% normal donkey serum (NDS) in PBS-T, then with primary antibodies (diluted in 3% NDS PBS-T) for 48hrs (4°C). Sections were then washed with PBS and incubated in the secondary antibodies for 2hrs at room temperature. A subset of sections was used to first validate the GFAP-GFP line i.e., verify GFP expression in GFAP+ astrocytes by double labelling for GFP (chicken anti-GFP, Aves GFP-1020, CA, USA followed by secondary donkey α-chicken AF488 antibody, Jackson Immuno Research 703-545-155, PA, USA) and antibodies against other cell types. The antibodies used were either the rabbit anti-GFAP antibody (Dako Z0334, CA, USA) followed by secondary antibody (donkey anti-rabbit DyLight405, Jackson Immuno Research 711-475-152), the ionized calcium binding adaptor molecule 1 (Iba1, microglial marker, rabbit anti-Iba1, Wako 019-19741, VA, USA) followed by donkey anti-rabbit DyLight405 (Jackson Immuno Research 711-475-152) or neuron nuclei protein marker (NeuN, neuronal marker, guinea pig anti-NeuN, EMD Millipore ABN90, ON, Canada) followed by donkey anti-guinea pig (CF405M, Biotium 20376, CA, USA).

### Acquisition and quantification

Using a confocal microscope (Olympus, Pennsylvania, USA) with oil-based 100X objective, 6 GFAP+ and 6 GFP+ cells selected from the gray matter, specifically from the anterior cingulate cortex region of the PFC of each mouse were imaged. To ensure the accurate Sholl analysis measurements, only non-overlapping immuno-positive cells were selected. The Z-axis diameter for each cell was determined by identifying the upper and lower Z-plane, and the middle of the astrocyte was set as Z0. The FluoView software (FV10-ASW 4.0b) was used to image five (GFAP+ cells) or three (GFP+ cells) z-stacks equally spanning Z0 (z-step = 5μm or 10μm, respectively). The z-steps for each cell type was optimized based on pilot acquisition and quantification trials for each staining. When using z-stacks of smaller intervals we found that the astrocyte morphology was not quantifiable and did not allow for distinguishing individual processes. The five (GFAP+ cells) or three (GFP+ cells) z-stacks Z-stack were projected on a bidimensional image upon which the Sholl analysis is performed using ImageJ’s plug-in FIJI (Maryland, USA) [63] and resulting images were converted to 8-bit, optimized for brightness-contrast before thresholding to minimize background noise **(Figure 2a-f)**. Images were then processed with a median filter (FIJI: despeckle function), which replaces each pixel with the median value in its 3 × 3 neighborhood removing any salt and pepper noise classically found after thresholding **(Supplementary Figure S4a-b)**.

**Figure 2:**
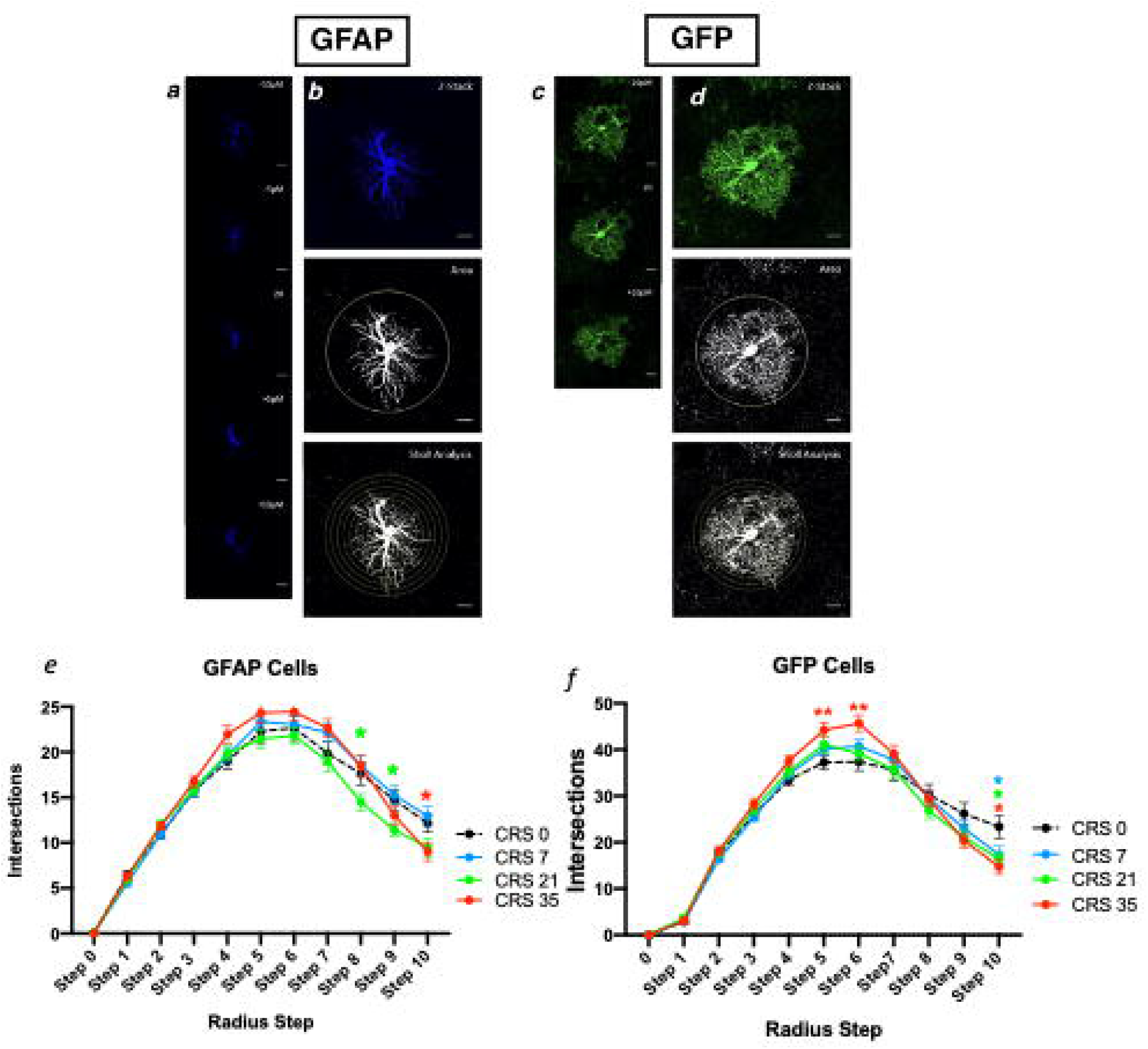
Chronic restraint stress (CRS) alters cortical astrocyte morphology. Scale = 10um. Representative astrocyte stained for glial fibrillary acidic protein (GFAP; blue) **(a**,**b**,**e)** or green fluorescent protein (GFP; green) **(c**,**d**,**f)**. Each GFAP stained cell is imaged over a span of 20μm, every 5μm, with the middle of the cell= Z0 **(a)**. For each GFAP cell, images are z-stacked **(b)**, thresholded and Sholl circles placed for analysis with the farthest circle placed at the end of the longest process. Each GFP stained cell is imaged over a span of 20μm, every 10μm, with the middle of the cell= Z0 **(c)**. For each GFP cell, images are z-stacked, thresholded and Sholl circles placed for analysis with the farthest circle placed at the end of the longest process **(d)**. CRS alters the number of intersections per radius step of GFAP+ cells **(e)** and GFP+ cells **(f)**. Data is presented as mean ± s.e.m. *\* p<0*.*05* and ** *p<0*.*01* compared to control group.

We first used the classical Sholl analysis methods with fixed radius steps of 5 or 10um as well as other parameters such as longest process, optical density over area and total number of intersections (see **Supplementary Material** for details). For the adapted Sholl analysis, the longest process of each GFAP+ and GFP+ cell was considered as the furthest Sholl circle (determined automatically in FIJI). Radius step was calculated as radius length divided by 11, obtaining a set number of 10 radius steps per cell (from 0-10). The number of intersections per radius step was defined as number of times processes cross each delineated circle and measured automatically.

### Statistical Analysis

For all behavioral data, ANOVA was used to determine the main effects of stress or sex for one-time readouts, and repeated measures ANOVA to identify potential interactions with time for the collected longitudinal data, followed by Fisher’s post-hoc test using StatView software 5.0 (CA, USA). For the Sholl analysis, power analysis performed on a previous published study [64] studying the effects of stress on amygdala and hippocampal astrocytes revealed that a minimum of 164 cells would be sufficient to detect an effect size of 0.3, with a 0.9 power and 0.05 probability. In our study, we quantified 6 astrocytes per marker per animal per group (192 cells). Repeated measure ANOVA was used to analyze the number of intersections per radius step. Finally, potential relationship between the complexity of cell morphology and behavior was determined using correlation analysis (Pearson’s R Test, Prism 8 Software, CA, USA).

## Results

### CRS mice exhibit poor coat state, anhedonia- and anxiety-like behaviors

Mice were exposed to 0, 7, 21 or 35 days of CRS and coat state, sucrose consumption and residual avoidance were measured weekly (**Figure 1a**). Weekly performances are reported in **Supplementary Material** and **Supplementary Figures S1 and S2**. Here, we present the data obtained in the last week of behavioral testing.

Analysis of coat state degradation scores revealed a significant main effect of stress (F_(3,28)_=18.349, p<0.0001). Mice subjected to 21 and 35 days of CRS had significantly increased coat state degradation compared to control mice (p<0.001 and p<0.0001 respectively, **Figure 1b**). Statistical differences between each group are reported in **Supplementary Table S1**. When sex was included as a factor, there was a main effect of sex (F_(1,28)_=22.059, p<0.0001; males have a higher overall score of coat state degradation than the females at baseline) but no sex*stress interaction **(Supplementary Table S1)**.

We found a significant main effect of stress on sucrose intake (F_(3,28)_ =3.952, p<0.05). Comparison between groups revealed significant sucrose intake reduction between mice exposed to 35 days of CRS compared to controls (p<0.01; **Figure 1c**), with no difference in water consumption between group (CRS0:1.4±0.2; CRS7:1.9±0.1; CRS21:1.7±0.2; CRS35:1.3±0.3). No significant effect of sex (F_(1,28)_=3.01, p=0.10) or a sex*stress interaction (F_(3,28)_=0.994, p=0.412) was found on sucrose intake **(Supplementary Table S1**).

Analysis of the time spent in the shelter zone calculated for five hours after the light challenge showed a significant stress effect (F_(3,28)_ =3.662, p<0.01). Post-hoc analysis revealed a significant increase in time spent in the shelter zone in all stress groups compared to the controls (p<0.01 for CRS 7 and p<0.05 for CRS 21 and 35; **Figure 1d**). When sex was considered as a factor a significant main effect of sex (F_(1,28)_=16.775, p<0.001) with males spend more time in the shelter than the females at baseline, but no stress*sex interaction (**Supplementary Table S1**). When looking at the hourly record of the animals’ time in the shelter zone, we found a significant main effect of time (F_(3,12)_=24.269, p<0.0001), stress (F_(3,27)_=3.757, p<0.05), but no significant interaction of stress over time (F_(3,36)_=0.979, p=0.508) (**Figure 1e**).

When considering Z-emotionality scores, we found a significant stress effect (F_(3.28)_ =16.451, p<0.0001) explained by a significant increase in the CRS 7, CRS 21 and CRS 35 groups compared to the controls (p<0.01, p<0.0001; **Figure 1f and Supplementary Table S1**). When sex was considered as a factor, a significant main effect of sex (F_(1,28)=_9.380, p<0.01) and a group*sex interaction (F_(3,28)_=13.136, p>0.0001) were found. Females showed greater increase in z-behavioral emotionality scores then males in response to chronic stress exposure (**Supplementary Table S1**).

### Verification of the GFAP-GFP Transgenic mouse line

Fluorescence immunohistochemistry and confocal analysis was performed on PFC sections from GFAP-GFP heterozygous mice to determine if GFP+ cells of this mouse line were co-labelled with either NeuN, Iba1 or GFAP staining. We found no overlap between GFP+ and NeuN+ cell populations. Similarly, GFP+ and Iba1+ cells represented two distinct cell types with no overlapping immunostaining. We qualitatively found that a vast majority of the GFP+ cells were GFAP co-labeled, validating the mouse line specificity. We decided to use both markers in our assessment of cell morphology as GFAP immunohistochemistry allowed for staining of primary processes while, GFP allows for detection of smaller processes (**Supplementary Figure S3**).

### CRS decreased the number of intersections at distal radius steps, suggesting atrophy of GFAP+ and GFP+ cells

We began our morphological analysis by immunolabelling and imaging sections from the PFC with labelled GFAP+ or GFP+ cells. We then thresholded the images for Sholl analysis as seen in **Figure 2a-f**. We were then able to analyze astrocytes using the classical Sholl technique with a fixed radius step of 5 or 10um (**Supplementary Figure S4c-e)**, longest process (**Supplementary Figure S4h-i**), optical density over area (**Supplementary Figure S4j-k**) and total intersection numbers (**Supplementary Figure S4l-m**). The results indicate no significant effect of CRS or sex (see **Supplementary Material** for details).

As astrocyte size and morphology heterogeneity may be a confounding factor when using the classical Sholl analysis, we adapted the method to quantify intersections per radius step (fixed number of radius steps = 11). For GFAP+ cells, there was a significant main effect of stress and radius step (F_(30,1880)_=2.468, p<0.0001; F_(3,10)_=379.990, p<0.0001; **Figure 2e**). The number of intersections for GFAP+ cells were significantly different at step 8, 9 and 10. Specifically, significant reduction in number of intersections were found between CRS0 and CRS21 at step 8 and 9, as well as between CRS 0 and CRS35 at step 10 (p<0.05; **Figure 2e)**. For GFP+ cells, there was also a significant main effect of stress and radius step (F_(30,1580)_=2.984, p<0.0001; F_(3,10)_=534.843, p<0.0001; **Figure 2f**). The number of intersections for GFP+ cells was significantly different between groups at step 5, 6 and 10. Specifically, CRS35 groups showed significantly increased at step 5 and 6, and significantly reduced at step 10 compared to CRS0. Similar reductions between CRS0 and CRS7 or CRS21 at step 10 were found (p<0.05; **Figure 2f**. In summary, at step 10, both GFAP+ and GFP+ cells show reduction of the number of intersections (F_(3,188)_=3.506, p<0.05; F_(3,158)_ =3.682, p<0.05; **Figure 3a-b**). ANOVA performed at this step radius also revealed a significant main effect of sex for GFP+ (F_(1,158)_=4.099, p<0.05) greater number of intersection in females compared to males) but not GFAP+ cells (F_(1,188)_=0.001, p=0.9840). For both cell types at this radius step, no group*sex interaction was found (F_(3,188)_=0.693, p=0.655; F_(1,158)=_1.599, p=0.192) (**Supplementary Table S1**).

**Figure 3:**
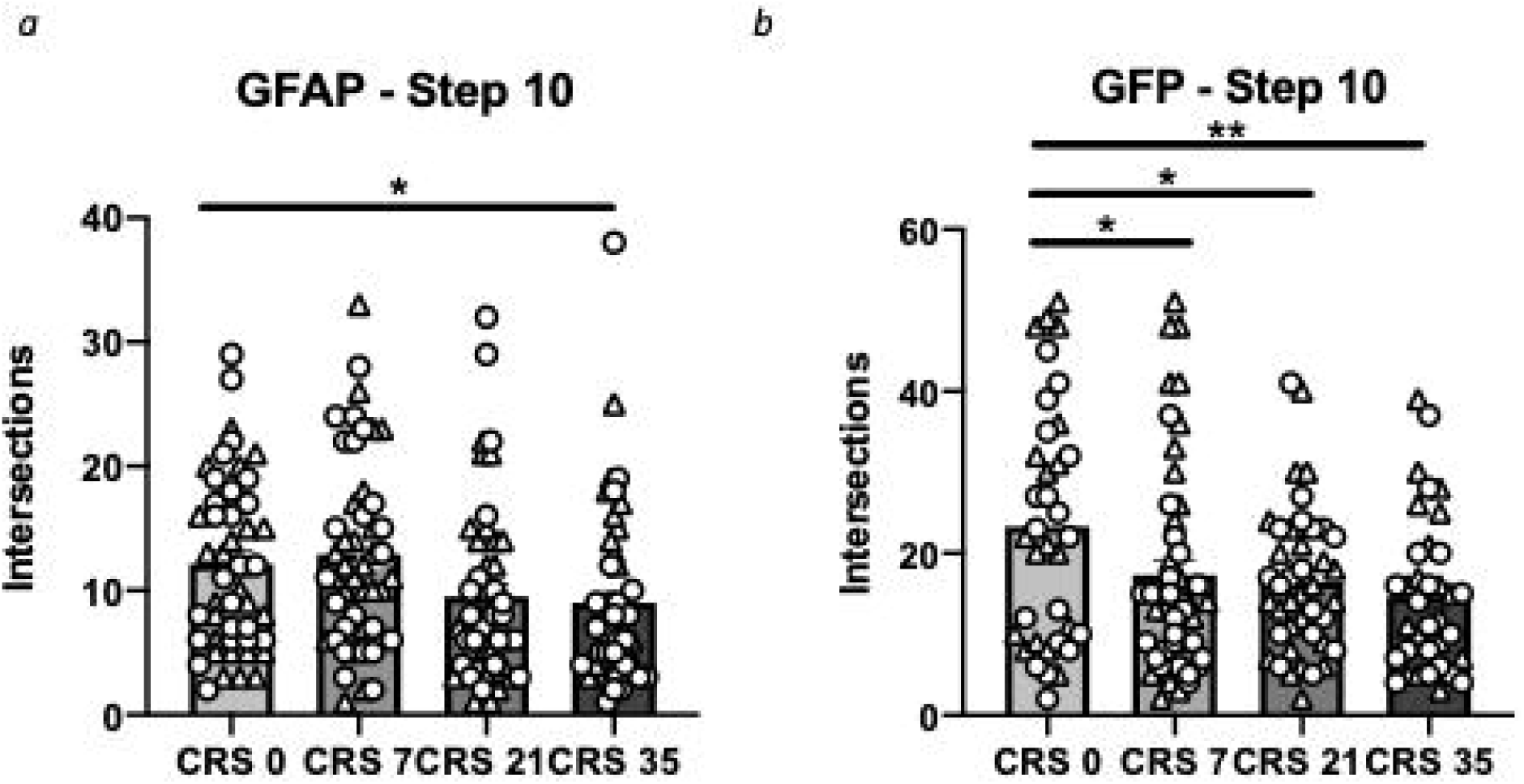
Chronic restraint stress (CRS) alters cortical astrocyte morphology at radius step 10. **(a)** CRS reduces the number of GFAP cell distal intersections at radius step 10. **(b)** CRS reduces the number of GFP cells distal intersections at radius step 10. Data is presented as mean ± s.e.m. *\* p<0*.*05* and ** *p<0*.*01* compared to control group. Females are denoted with a triangle and males with a circle.

### CRS-induced reduction in the number of intersections at distal radius steps correlates with anhedonia-like behavior and with z-behavioral emotionality scores

Potential correlations between anhedonia-like behavior (sucrose intake) or anxiety-like behavior (shelter time) and altered astrocyte morphology were then investigated. We found no correlation between radius step 10 intersections and sucrose intake (*r*=0.233, p=0.200; **Figure 4a)** or time in the shelter (*r*=0.107, p=0.559, **Figure 4c**) for GFAP+ cells. No correlation was found when the analysis was performed separately for males (sucrose intake: *r*=0.357, p=0.175; time in shelter: *r*=-0.159, p=0.556) and females (sucrose intake: *r*=0.003, p=0.992; time in shelter: *r*=0.410, p=0.115). There was a positive correlation between radius step 10 intersections and sucrose intake for GFP+ cells (*r*=0.489, p<0.01; **Figure 4b**). This positive correlation with GFP+ cells was also significant when data was analysed separately for males (*r*=0.622, p<0.05) and marginally significant for females (*r*=0.547, p=0.053). In addition, there was no significant correlation between the number of intersections at step 10 and time spent in the shelter for GFP+ cells overall (*r*=-0.248, p=0.212, **Figure 4d**) and in females (*r*=-0.253, p=0.344). However, in male mice there is a trend towards a significant negative correlation (*r=*-0.517, p=0.070). Finally, there was a trend towards a significant negative correlation between z-emotionality score and number of intersections at step 10 for GFP+ (*r*=-0.341, p=0.08; **Figure 4f**) but not for GFAP+ cells (*r*=-0.257, p=0.156; **Figure 4e**). In GFP+ cells, male mice showed a significant negative correlation between number of intersections at radius step 10 and z-emotionality score (*r*=-0.667, p<0.05) but not female mice (*r*=-0.340, p=0.235). GFAP+ intersections at radius step 10 did not correlate with z-emotionality in males (*r*=-0.411, p=0.114) or female (*r*=-0.235, p=0.344). Due to the discrete changes in astrocyte morphology, the significant correlations between number of intersections and behavioral performances did not survived correction for multiple comparisons. However, we found similar correlation levels at radius step 9 (**Supplementary Figure S5**).

**Figure 4:**
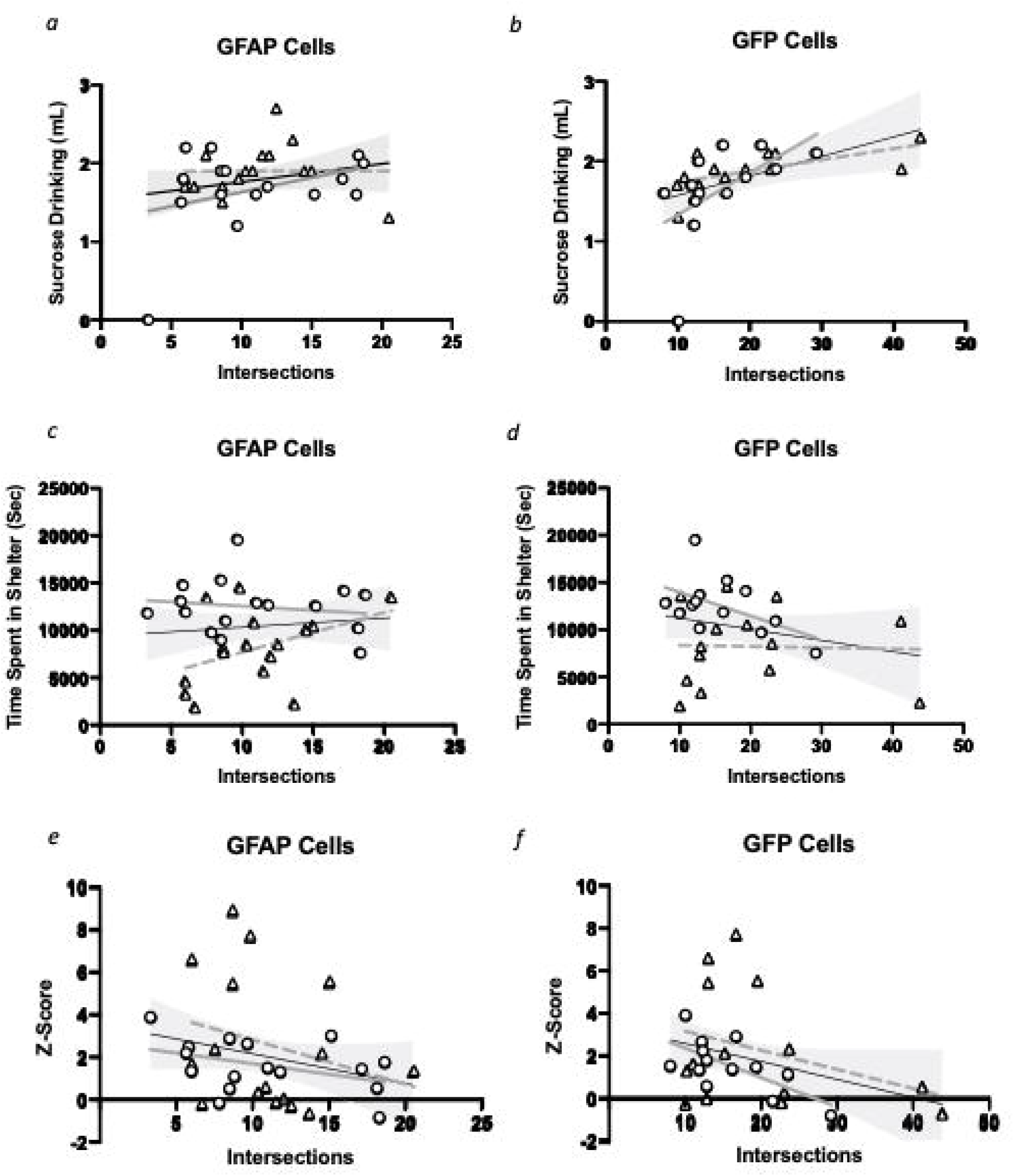
Chronic restraint stress (CRS)-induced alterations in cortical astrocyte morphology correlates with behavioral performances. The average number of intersections at step 10 for each animal calculated for glial fibrillary acidic protein (GFAP) **(a**,**c**,**e)** or green fluorescent protein (GFP) positive cells **(b**,**d**,**f)** (**Supplementary Figure 4)**. Relationship between GFAP-cell intersections at step 10 and sucrose intake **(a)**, time spent in the shelter **(c)** and z-score behavioral emotionality **e)** was assessed using Pearson’s correlational analysis. Overall regression lines are black. Females are denoted with a triangle, with grey dotted regression lines and males with a circle, with grey bolded regression lines. **(a)** Overall: *r*=0.099, p=0.517; Females: *r*=0.003, p=0.992; Males: *r*=0.357, p=0.175, **(c)** Overall: *r*=0.107, p=0.559; Females: *r*=0.410, p=0.115; Males: *r*=-0.159, p=0.175, **(e)** Overall: *r*=-0.257, p=0.156. Females: *r*=-0.253, p=0.344; Males: *r*=-0.411, p=0.114. A similar analysis was performed for GFP-cell intersections at radius step 10 and sucrose intake **(b)**, time spent in the shelter zone **(d)** and z-score behavioral emotionality **(f). (b)** Overall: *r*=0.489, p<0.01; Females: *r*=0.622, p<0.05; Males: *r*=-0.547, p=0.053, **(d)** Overall: *r*=-0.248, p=0.212; Females: *r*=-0.032, p=0.914; Males: *r*=-0.517, p=0.070, **(f)** Overall: *r*=-0.341, p=0.082; Females: *r*=-0.340, p=0.235; Males: *r*=-0.667, p=0.013. Grey area represents 95% confidence intervals.

## Discussion

In this study, we investigated the astroglial morphological changes induced by CRS in the mouse PFC. We first confirmed that CRS-exposed mice displayed anxiety- and anhedonia-like behaviors. Indeed, CRS-exposed mice showed increase in time spent in the shelter zone in the Phenotyper test after 7, 21, and 35 days of CRS. CRS also induced anhedonia-like behavior i.e., decreased sucrose intake following 35 days of CRS. When considering the average Z-behavioral emotionality score of all behavioral tests, CRS increased behavioral emotionality following 21 and 35 days of CRS. At the cellular level, we improved upon current methods of Sholl analysis to account for intrinsic heterogeneity of astrocyte size [54], and found that CRS reduced the number of intersections at distal radius steps of GFAP+ cells following 21 and 35 days of exposure. Similar CRS-induced decreases in intersections at distal radii for GFP+ cells were found, accompanied by increased proximal processes, suggesting atrophy of the GFAP+ and GFP+ cells. For PFC GFP+ cells, distal process intersection reductions that correlated with sucrose intake (in both males and females), but not with time spent in the shelter zone Interestingly, for GFP+ cells, the reductions in the distal intersections trended towards a negative correlation with greater z-emotionality, an effect not seen in GFAP+ cells. GFP+ cell atrophy was more pronounced and significantly negatively correlated with z-score in male mice. Altogether, our results showed alterations of astrocyte morphology linked to the behavioral response to CRS. This suggests that morphological reorganization of astrocytes and atrophy may contribute to the behavioral deficits induced by chronic stress.

The CRS paradigm is a well-documented rodent model of chronic stress that was shown to alter the physical animal’s state and produce behavioral deficits, including anxiety- and anhedonia-like behaviors [65-69]. Here, we confirmed previous findings from our lab and others demonstrating greater coat state deterioration [60, 61], reduction in sucrose intake [69] and increase in time spent in the shelter zone after the light challenge in the PhenoTyper test following CRS [61]. Importantly, we opted for the use of the CRS paradigm, instead of the chronic unpredictable (mild) stress model which employs milder, randomized, and variable stressors, presuming that CRS would be ideal for a time course study, as it would allow for detection of more distinct behavioral and cellular differences between groups with increasing exposure to stress [68, 70]. In addition, the CRS paradigm was widely used to examine neuronal morphology changes in the PFC, hippocampus, and amygdala [2, 44, 70-73].

The majority of studies investigating neuronal morphology classically use the Sholl analysis which consists of placing concentric circle (or spheres) on 2-D images (or 3-D images) of neurons, extending from the soma to the tip of the apical dendrite. Each circle is placed at equal distance i.e. fixed radius step size (commonly 10-20 µm), constant regardless of the cell size [47]. The apical dendrite length is often selected as a scalar value of complexity to be employed in follow-up correlative analysis within an experiment [47, 70, 74]. This approach has been extensively and successfully used to report the neuronal morphological alterations associated with chronic stress [70, 75, 76]. We began our investigation by applying these parameters to analyze GFAP+ and GFP+ cell morphology but were unable to detect differences between groups. In this scenario, the use of a fixed radius step, with a final Sholl circle placed at the same distance for small or larger astrocyte of similar complexity, would skew the data as the smaller astrocyte would artificially display no quantifiable intersections at the farthest Sholl circle (**Supplementary Figure S6**). This highlights the need to adapt the Sholl analysis method to astrocytes to account for the size heterogeneity of this cell population [54]. Investigators have attempted to address these methodological challenges using parameters such as cell perimeter [77], soma size or volume. Chan et al. [52] investigated CRS effects on GFAP+ soma size and volume but were unable to find significant changes on either parameter. Other alternate methods were used to complement the Sholl analysis to address differences in morphological complexity of astrocytes. For example, Johnson et al., [78] employed a parallel qualitative manual estimation to separate the cells into complexity groups ; however, the complexity assessment was subjective, even if the analysis was performed by a blinded experimenter [78]. In the present study, to ensure that the distance between Sholl circles is dependent on the size of the cell and that equivalent samples are quantified for each astrocyte, we incorporated cell size in the analysis by using a set number of radius steps. The results of this method unveiled reductions of intersection number at the distal Sholl circles in mice subjected to CRS whether the cell was GFAP or GFP positive. GFP+ cells of CRS animals also showed an increase in number of intersections at the proximal Sholl circles. This data demonstrates reorganisation of astrocyte morphology following CRS, and suggest distal process atrophy along with preservation of processes closer to the soma. Distal process loss was marked at radius step 10 for both GFAP+ and GFP+ cells. CRS effects on astrocyte morphology appeared earlier and of greater magnitude when quantifying GFP+ cells. This may not be surprising since CRS may affect fine astrocytic processes first (GFP stained processes) and primary (GFAP stained) processes later. Ultimately, we adapted the Sholl analysis to allow for a more refined assessment of astrocyte morphology and demonstrated an atrophy of astrocyte distal processes following CRS.

To establish a relationship between behavior and PFC astrocyte morphology, we investigated whether intersections at distal radius step 10 correlated with anhedonia- and anxiety-like behaviors. We found that greater astrocyte atrophy at this radius step positively correlated with greater anhedonia-like behavior, but not only trended with greater anxiety-like behavior. These correlations were more pronounced in males. Overall, we also demonstrated a link between GFP+ cell morphology and Z-behavioral emotionality score, suggesting that PFC astrocyte alterations following CRS contribute to behavioral emotionality. Our findings are consistent with the data of the literature showing that chronic stress reduced PFC astrocyte number or function [39, 79] and that ablation or dysfunction of these cells induce depressive-like behaviors [32, 38, 80]. Similar astrocytic loss and dysfunction were also reported in other brain regions and were involved in the expression of emotion-related deficits [2, 37, 40]. In addition, a recent study demonstrated that astroglial morphology is altered in specific subfields of the hippocampal formation following chronic mild unpredictable stress, however no correlation with behavioral outcomes was performed [81]. These additional effects of chronic stress on astrocytic loss or dysfunction as well as the involvement of astrocytic changes in others brain regions may account for additional astroglial contribution to behavioral emotionality.

The chronic stress-induced PFC astroglia atrophy identified in this study may supplement the already well-established neuroplastic changes associated with chronic stress [1, 2] or MDD [14, 19, 20, 82]. It is possible that the same mechanisms involved in the chronic stress-induced neuronal atrophy also alters astrocyte morphology e.g. increased corticosterone [3, 83], excess extra-synaptic glutamate [4] or decreased trophic support [71]. Astrocytes also play a vital role in the regulation and function of neurons, particularly through the reuptake of extra-synaptic neurotransmitters [23, 26, 27, 32]. Therefore, chronic stress-induced alterations of astrocyte morphology could have direct consequences on the neuronal function and synaptic transmission, exacerbating chronic stress effects on neuronal populations. To investigate this hypothesis further, functional consequences of manipulations mimicking chronic stress effects on astrocyte morphology, activity or function should be considered. These studies may include investigating activity or cell signaling changes within astrocytes after chronic stress or neurons following astrocyte manipulations [69, 84-87].

Although the main focus of this study was investigating the effects of chronic stress exposure on astrocyte morphology, according to current research guidelines, we included an equal number of male and female mice in our study design to account for sex a basic biological variable. Interestingly, CRS male mice showed greater anxiety- and anhedonia-like behavior, as well as increased degradation of coat state when individual behavioral outcomes are considered. However, CRS females showed overall greater Z-emotionality score normalized to controls of the same sex group across behavioral tests. This is consistent with literature suggesting that females are more effected by stress and exhibit more and greater symptoms in stress-related disorders [88]. Although we confirmed sex differences in the behavioral response to CRS [61, 69], we only found some minor sex differences in morphology, and we were unable to see interaction between sex and groups. This could indicate that chronic stress does not alter astrocyte morphology in a sex-dependent manner or that considerably greater number of animals per group and per sex are need to detect sexual dimorphism associated with PFC astrocyte morphology, as previously described in the hypothalamus [89]. Future studies specifically designed for studying sexual dimorphism of astrocyte morphology in PFC would be needed to answer this question. These studies may also need to include estrus cycle stages as an additional variable since this may explain why similar but not significant correlations between PFC astrocyte morphology and behavioral changes were found in males and females.

This study is not without its limitations. First, as mentioned above, we conducted the study on relatively small number of animals of each sex. Second, we chose to focus on GFAP+ astrocytes and not S100β+ cells, as mice exposed to chronic stress show only reductions in GFAP+ cells [40]. We also focused on the PFC as this is the main region consistently showing astroglial changes in cell density in MDD. Findings in other brain regions such as the hippocampus have been inconsistent [81, 90, 91]. Finally, this study focused on identifying chronic stress effects on PFC astrocyte morphology; the mechanisms responsible for chronic stress-induced atrophy or the consequences of these changes on neuronal morphology, activity and function remain to be investigated. Indeed, these astrocytic changes may be linked to the decrease in synapse number and synaptic markers reported in chronic stress models [2, 92] and MDD [18, 19, 93].

Nevertheless, in this study we demonstrated that chronic stress alters GFAP+ astrocyte morphology in the PFC and may contribute to the chronic stress-associated behavioral deficits. We examined the timeline of these effects and provided a quantitative method of analysis of astrocyte complexity. This complements preclinical studies that consistently show reduced density and number of GFAP+ astroglia cells in chronic stress models and findings of astrocyte reduction in MDD further highlighting the role of astrocytes in stress-related illnesses.

## Supporting information

Supplementary Section

## Acknowledgements

We would like to thank Colin Arrowsmith for his input on the calculations of astrocyte morphology indexes. We would also like to acknowledge the contributions of the CAMH animal facility personnel for animal care and genotyping services, specifically: Lori Dixon, Kristen Fournier, Katrina Deverell, and German Fernandes.

## Significance Statement

Astrocytes are brain cells pivotal to the normal function and support of neurons. Clinical and preclinical studies have consistently shown astroglial anomalies in depression and chronic stress rodent models. Here we demonstrate that chronic stress in mice induces atrophy of the prefrontal cortex astrocytes. We also found that these morphological changes correlate with the behavioral deficits associated with chronic stress. Altogether, our results strongly suggests that stress-induced morphological reorganization of astrocytes and atrophy may contribute in part to the behavioral deficits induced by chronic stress. This work further highlights the contribution of astroglial pathology in stress-related disorders, including depression.

