## Supplementary Section for "Chronic Stress Alters Astrocyte Morphology in Mouse Prefrontal Cortex"

**Supplementary Methods:**

*Removing Background before morphology analysis:*

As mentioned in the main text, when images of the astrocytes are thresholded, a significant background remains. **Supplementary Figure S3a** illustrates a representative image before thresholding. The salt and pepper background may impair the ability to confidently quantify the number of intersections of the Sholl circles, as a random “speck” over the Sholl circle could be counted as an intersection. Therefore, to reduce the number of false positive intersections, we used the Fiji despeckling tool in the ImageJ Fiji package (Maryland, USA). The principle of despeckling consists in applying a median filter based on pixel color. Briefly, it works by replacing each pixel with the median value in its 3 x 3 neighborhood. In other words, for each pixel, the center pixel replaced with the median value by the nine pixels surrounding pixel. This means that all lonely outliers are removed **(Supplementary Figure S3b)**. This method of background removal is not perfect as it is possible that some background could be left over, but using the Fiji despeckling tool multiple times, although possible, significantly reduces signal. To confirm this technique, we compared p-value results of images that were despeckled to ones that were not and, found that the results were remarkably close. Using the Fiji despeckling tool once, slightly strengthened and weakened some effects, as to be expected, but did not drastically change any results (or make anything significant that was not already).

*Employing classical Sholl analysis on astrocyte vs. adapted methodology:*

We first attempted to quantify astroglial morphological changes using the Sholl analysis as classically employed for neurons: placing concentric circle (or spheres) on 2-D images (or 3-D images) extending from the soma to the tip of the longest processes (or apical dendrite for neurons). Each circle was placed at equal distance i.e. fixed radius step size (commonly 10-20 µm) that is constant regardless of the cell size. In this study, we used 5 µm or 10 µm step sizes as the astrocytes were generally smaller (**Supplementary Figure S4c-e)**. This method was not ideal for analysis with astrocytes as they are highly heterogeneous in size which can lead to the problem we display in **Supplementary Figure S6**. In **Supplementary Figure S6a,** we show a schematic representation of an astrocyte with Sholl circles placed at fixed radius step distances (e.g 5µm steps) and the related number of intersections quantification **(Supplementary Figure S6b)**. However, when the same astrocyte is represented smaller in **(Supplementary Figure S6c)** with the same branching complexity, the analysis of the number of intersections reveals a relatively low number of branching crossing the distal circles and overall considerably lower total number of intersections **(Supplementary Figure S6d)**. Indeed, in this scenario, the use of a fixed radius step, with a final Sholl circle placed at the same distance for either astrocyte, bias the data, as the smaller astrocyte would artificially display no quantifiable intersections at the farthest Sholl circle. In order to adapt this method to better reflect astrocyte morphology, we use a distance between Sholl circles that is dependent on the size of the cell, by utilizing a set number of radius steps allowing for equivalent sample quantified for each astrocyte (**Supplementary Figure S6e-f)**. Data obtained using this methodology can be found in in the main manuscript.

Changes in other morphological parameters or measures commonly used for neurons were also quantified. We quantified the length of the longest process to attempt to recreate the common neuronal measure of apical length (**Supplementary Figure S4f-g).** Note that this approach was not an ideal as most astrocyte processes are very similar in length. We also quantified the intensity of the GFAP or GFP signal over the area covered by the astrocyte, through measuring optical density (OD) and using the area of the outer Sholl circle (**Supplementary Figure S4h-i)**. We also quantify the total intersections (**Supplementary Figure S4j-k)** and the average of intersections per animal at distal radius step 10 or both GFAP+ and GFP+ cells for use in correlational analysis (**Supplementary Figure S4l-m)**. Finally, in order to look more closely at the correlation of morphology and behaviour, we correlated intersections at radius step 9 with the behavioural task in the same manner as radius step 10 in the main text (**Supplementary Figure S5**). These correlations were also done using Pearson’s R test.

**Supplementary Results:**

*Time lapse of behavioural effects reveals changes in groups during longitudinal assessment of chronic restraint stress:*

Behavioral data of PhenoTyper tests for shelter zone across was assessed various time points (every week) throughout the 35 days the experiment. We used repeated-measures ANOVAs to determine if there is a significant difference in the time spent in the shelter zone per hour per group at each time point. At baseline, there was no significant difference between the groups (F_(36.336)_=1.919, p=0.149; **Supplementary Figure S2a**). No significant difference between groups was detectable on week 1 (F_(36.336)_=1.853, p= 0.161; **Supplementary Figure S2b**),week 2 (F_(36.336)_=0.765, p= 0.523; **Supplementary Figure S2c**) or week 3 (F_(36.336)_=1.235, p=0.316; **Supplementary Figure S2d**). On week 4 and week 5, the repeated-measures ANOVA of time spent in the shelter was unable to detect a main effect (F_(36.336)_=1.919, p=0.149 and subsequently F_(36.336)_=0.925, p=0.596; **Supplementary Figure S2e-f**), probably because 3 of the 4 groups showed similar responses. Indeed, all 3 CRS animal groups spent significantly more time in the shelter after the light challenge compared to controls (**Supplementary Figure S2e-f)**.

Coat state degradation and sucrose drinking was also assessed weekly. Repeated measures ANOVA of coat state degradation revealed a significant effect of CRS over time (F_(15,140)_=16.28, p<0.0001; **Supplementary Figure S3a**). Post-hoc analysis a significant increase in degradation of the coat state of the CRS 35 animals at 2, 3, 4 and 5 weeks compared to controls. There was also a significant degradation of coat state of the CRS 21 animals at week 4 and 5 of testing (i.e. week 2 and 3 of CRS exposure) compared to the controls (**Supplementary Figure S3a)**. Longitudinal analysis of sucrose drinking or time in the shelter following the light challenge showed no significant main effect of stress (F_(15,140)_=1.572, p=0.089 and F_(15,140)_=1.876, p=0.157, respectively; **Supplementary Figure S3b-c**). No effect of stress was found on water consumption measured every week (week 1: F_(3,28)_=0.275, week 2; F_(3,28)_=0.956, week 3: F_(3,28)_=1.385, week 4: F_(3,28)_=0.151 and week 5: ; F_(3,28)_=2.657, all p>0.05).

*Supplementary Morphology Results:*

The results obtained by employing the classical Sholl analysis to astrocytes using a set radius step length of 5 μm or 10 μm revealed no effects of CRS exposure**.** Indeed, the analysis of the number of interactions / radius step (10 μm) for GFAP cells showed no significant main effect of stress (F_(3.188)_=0.885, p=0.450; **Supplementary Figure S4e**). Similarly, when a radius step size of 5 μm was used, we found no significant differences between groups for neither GFAP+ cells (F_(3.188)_=0.573, p=0.633; **Supplementary Figure S4c**) nor GFP+ cells (F_(3.158)_=1.243, p=0.296; **Supplementary Figure S4d**).

Effects of CRS were also investigated on other parameters of the Sholl analysis. We found no significant effects of CRS on the length of longest process (GFAP F_(3,188)_=0.051, p=2.632, GFP F_(3,158)_=1.809, p=0.148; **Supplementary Figure S4f-g**) or OD/ Area (GFAP F_(3,158)_=1.852, p=0.139, GFP F_(3.158)_=0.866, p=0.460; **Supplementary Figure S4h-i**).

We then adapted Sholl analysis using a fixed number of radius step (n=11) for each astrocyte. The results of the quantification of the average of the numbers of intersections per radius steps for each group can be found in the main manuscript (**Figure 2g-h**). The total number of intersections for the cells was also analyzed and there was no change (GFAP F_(3.188)_=1.858, p=0.138, GFP F_(3.158)_=0.511, p=0.675; **Supplementary Figure S4j-k**).

The average of number of intersections at distal radius step 10 was calculated per mouse. We found no significant effect of CRS on this astrocyte morphology parameter (GFAP+ cells: F_(3,188)_=1.519, p=0.214, GFP+ cells; F_(3,158)_=1.098, p=0.370; **Supplementary Figure S4l-m)**. The average number of intersections at distal radius step 10 per mouse was used for the correlation analysis of astrocyte morphology and behavioral performances (**Figure 4a-f**).

Finally, to verify the validity of the correlation observed at radius step 10, we also correlated the number of intersections at radius step 9 with the behavioural performances (**Supplementary Figure S5**). For GFAP+ cells, we found that the average number of intersections at radius step 9 did not significantly correlate with sucrose drinking (r=0.242, p=0.182) or time spent in the shelter (r=0.091, p=0.620). For GFP+ cells, the average number of intersections at radius step 9 did positively correlate with sucrose intake (r=0.481, p<0.01) but did not significantly correlated with time spent in the shelter in PhenoTyper test (r=-0.203, p=0.309). For both GFAP+ and GFP+ cells, did not see any correlations between the average number of intersections at radius step 9 and z-behavioural emotionality score (GFAP: r=-0.176, p=0.335; GFP: r=-0.269, p=0.175). These results follow a similar pattern as the finding observed at radius step 10 (**Figure 4a-f**).

**Supplementary Figures:**

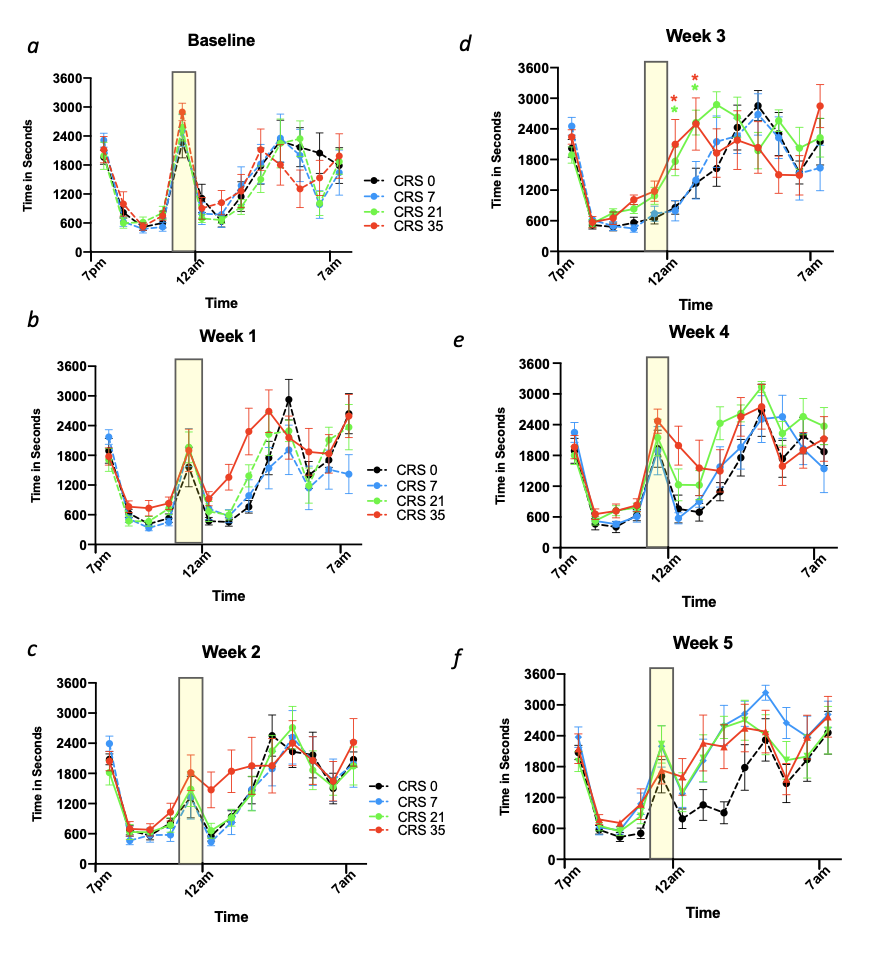

***Supplemental Figure S1: Time in the shelter zone hourly performances at baseline and each following week.*** Time spent in the shelter zone was monitored hourly from 7pm to 7am a baseline **(a)**, on week 1 **(b),** week 2 **(c),** week 3 **(d),** week 4 **(e)** and week 5 **(f)** of the experiment. Dashed line indicates when the animal group has not yet began CRS and the yellow box indicates the time the light is on. Data is presented as mean ± s.e.m.** p<0.05* compared to control group.

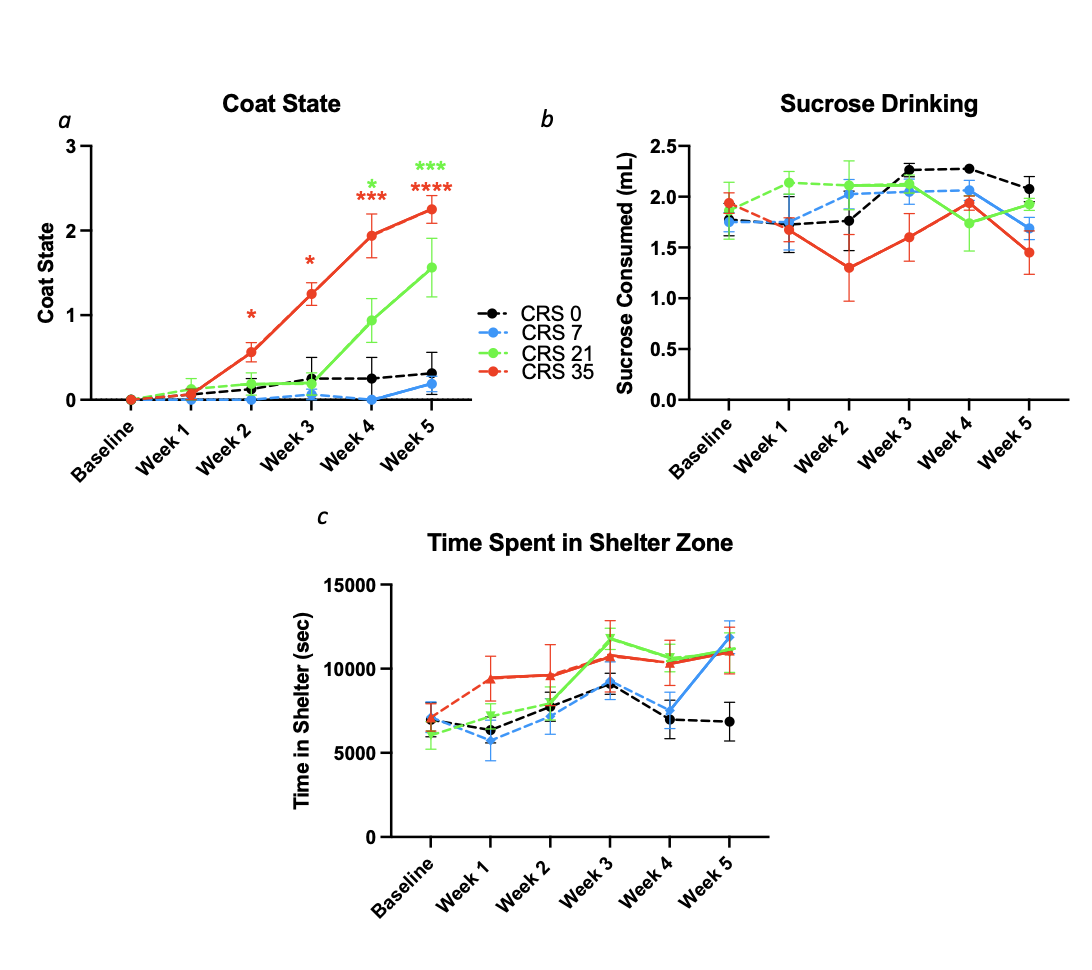

***Supplemental Figure S2: Trajectory of coat state, sucrose drinking, and time spent in the shelter zone at baseline and every following week.*** Coat state **(a),** sucrose drinking **(b)** and time spent in the shelter zone **(c)** was measured every week of the CRS experiment. Data is presented as mean ± s.e.m. Dashed line indicates the animal group has not yet began CRS.** p<0.05*, ***p<0.01*,and ****p<0.001, and ****p<0.0001* compared to control group.

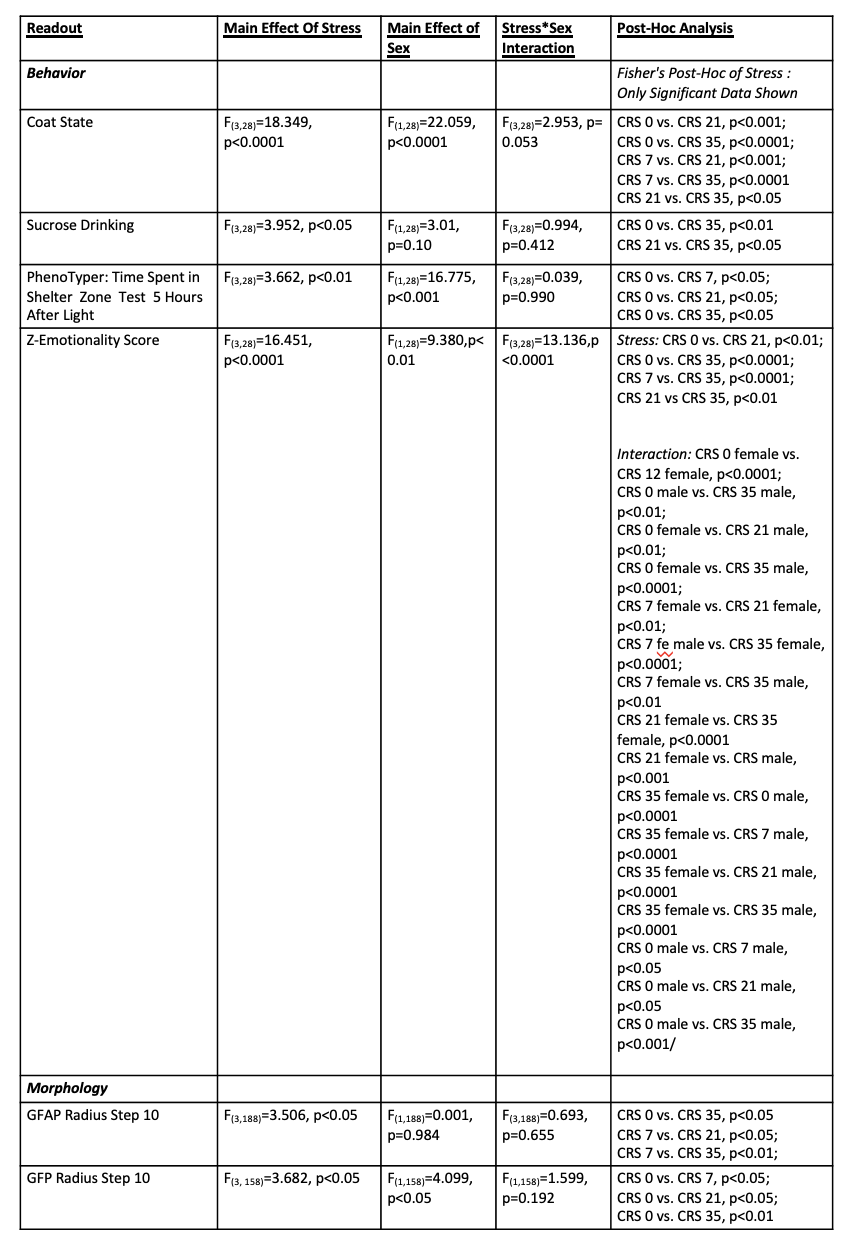

***Supplementary Table S1: Behavioural and Morphological statistical analysis with sex as a factor.***

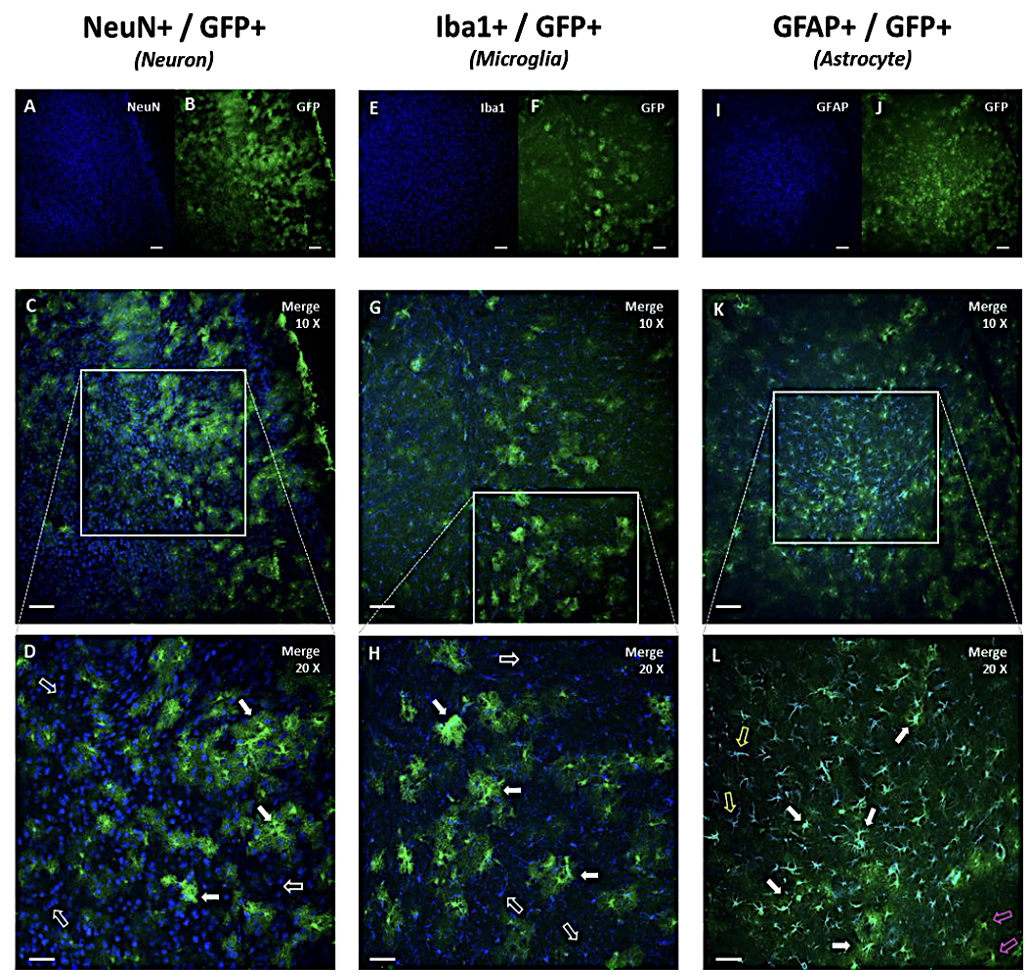

***Supplementary Figure S3: Representative images of GFAP-GFP mouse prefrontal cortex sections stained for GFP and, neuronal, microglia, or astrocyte marker.* (a-d)** Tissue immune-labeled for neuronal nuclei (NeuN) and green fluorescent protein (GFP). The filled arrows point at GFP+ cells, whilst the empty arrows indicating NeuN+ cells. **(e-h)** Tissue immune-labeled for ionized calcium binding adaptor molecule 1 (Iba1) and GFP. Filled arrows point at GFP+ cells and the empty arrows indicating Iba1+ cells. ***(i-l)*** Tissue immuno-labeled for glial fibrillary acidic protein (GFAP) and GFP. These filled arrows indicate GFAP+/GFP+ overlapping cells, while empty yellow arrows indicating only GFAP+ cells (GFAP+/GFP-), and empty pink arrows indicating only GFP+ cells (GFAP-/GFP+). Note that lack of perfect overlap between GFP+ and GFAP+ is most probably due to immunohistochemistry protocol conditions. Indeed, antibody concentrations were low and optimized for signal/noise ratio optimal for the morphological analysis. Scale bar represents 50um (d, h and l) and 100um of all other images.

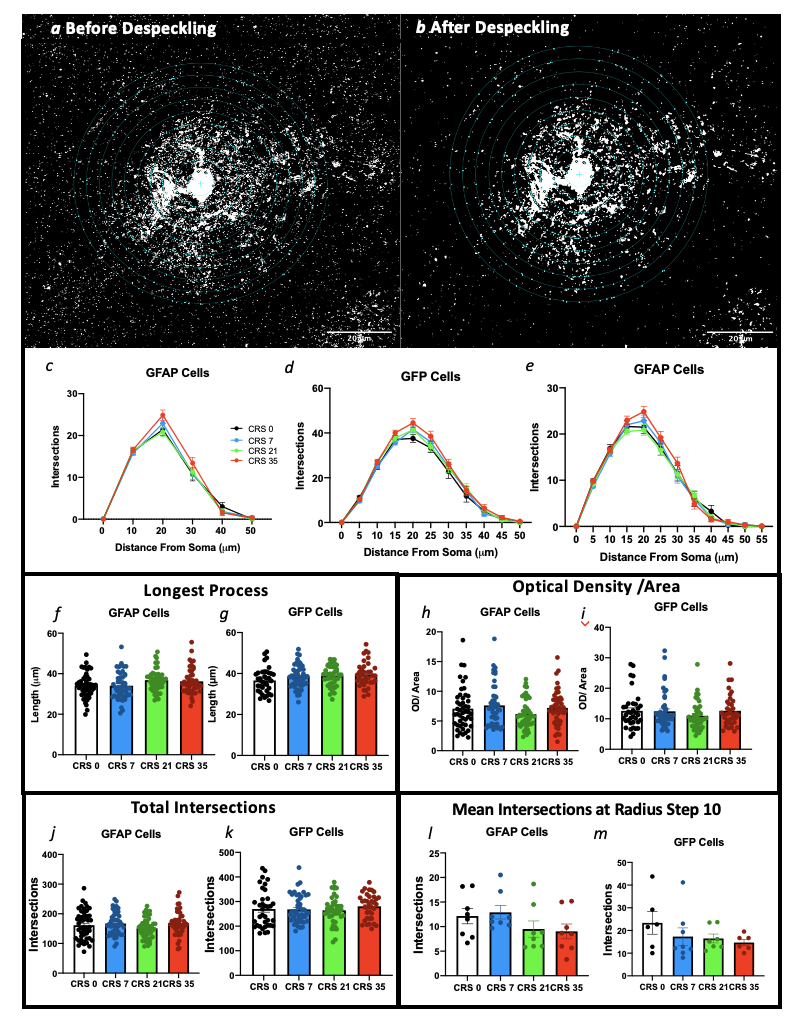

***Supplementary Figure S4: Sholl analysis methodology and additional data. (a-b)*** Representative images of an astrocyte visualized before and after despeckling. Analysis of number of intersections for glial fibrillary acidic protein (GFAP) cells using 10 um **(c)** or 5 um **(d)** radius step size. Analysis of the number of intersections for green fluorescent protein (GFP) cells using 5um radius step size **(e)**. Effects of CRS on the longest process of GFAP+ **(f)** and GFP+ cells **(g)**. Effects of CRS on optical density / area of GFAP+ **(h)** and GFP+ cells **(i)**. Effects of CRS on the total number of intersections for GFAP+ **(j)** and GFP+ cells **(k)**. Effects of CRS on distal intersections at radius step 10 averaged per mouse for GFAP+ **(l)** and GFP+ cells **(m)**.

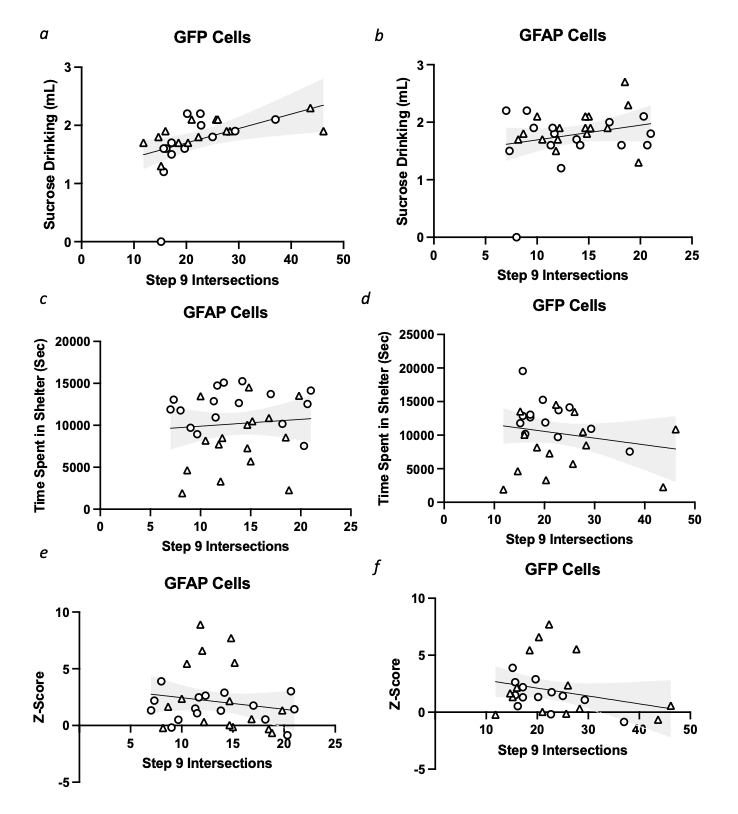

***Supplementary Figure S5: Chronic restraint stress (CRS)-induced alterations in cortical astrocyte morphology correlates with behavioral performances.*** The average number of intersections at step 9 for each animal calculated for glial fibrillary acidic protein (GFAP) **(a,c,e)** or green fluorescent protein (GFP) positive cells **(b,d,f)** (**Supplementary Figure 4).** Relationship between GFAP-cell intersections at step 9 and sucrose intake **(a)**, time spent in the shelter **(c)** and z-score behavioral emotionality (**e)** was assessed using Pearson’s correlational analysis. **(a)** r=0.242, p=0.182; **(c)** r=0.091, p=0.**620; (e)** r=-0.176, p=0.335. A similar analysis was performed for GFP-cell intersections at radius step 9 and sucrose intake **(b),** time spent in the shelter zone **(d)** and z-score behavioral emotionality **(f). (b)** r=0.481, p<0.0; **(d)** r=-0.203, p=0.309; **(f)** r=-0.269, p=0.175. The grey area represents 95% confidence intervals. Females are denoted with a triangle and males with a circle.

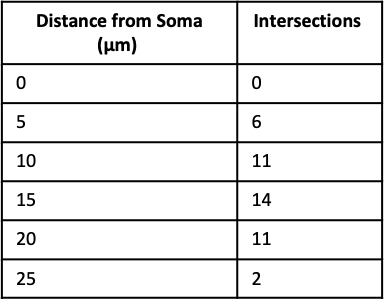

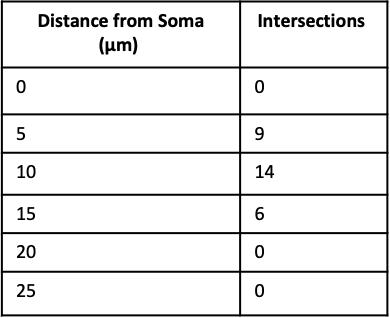

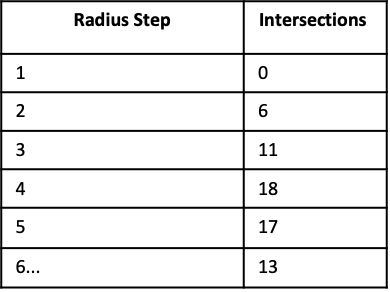

*b*

*a*

*c*

*d*

*e*

*f*

***Supplementary Figure S6: Schematic representation of two astrocytes of similar branching complexity but different size analysed using the two Sholl methodologies.*** ***(a-b)*** Illustration of an astrocyte quantified using Sholl circles placed at fixed radius step distances and related number of intersections quantification. ***(c-d)*** Illustration of same astrocyte but smaller quantified using Sholl circles placed at fixed radius step distances and related number of intersections quantification. ***(c-d)*** Illustration of same astrocyte but smaller quantified with a fixed number of radius step (e.g. 11) and related number of intersections quantification.
